# Glucose-fed microbiota alters intestinal epithelium and increases susceptibility to bacterial pathogens

**DOI:** 10.1101/2023.11.19.567723

**Authors:** Samuel F. Kingsley, Yonghak Seo, Alicia Wood, Khursheed A. Wani, Xavier Gonzalez, Javier Irazoqui, Steven E. Finkel, Heidi A. Tissenbaum

## Abstract

Overconsumption of dietary sugar can lead to many negative health effects including the development of Type 2 diabetes, metabolic syndrome, cardiovascular disease, and neurodegenerative disorders. Recently, the human intestinal microbiota strongly associated with our overall health has also been known to be affected by diet. However, mechanistic insight into the importance of the human intestinal microbiota and the effects of chronic sugar ingestion has not been possible largely due to the complexity of the human microbiome which contains hundreds of types of organisms. Here, we use an interspecies *C. elegans*-*E. coli* system, where *E. coli* are subjected to high sugar, then consumed by the bacterivore host *C. elegans* to become the microbiota. This glucose-fed microbiota results in a significant lifespan reduction accompanied by reduced healthspan including locomotion, stress resistance, and changes in behavior and feeding. Lifespan reduction is also accompanied by two potential major contributors: increased intestinal bacterial density and increased reactive oxygen species. The glucose-fed microbiota accelerated the age-related development of intestinal cell permeability, intestinal distention, and dysregulation of immune effectors. Ultimately, the changes in the intestinal epithelium due to aging with the glucose fed microbiota results in increased susceptibility to multiple bacterial pathogens. Taken together, our data reveal that chronic ingestion of sugar such as a western diet has profound health effects on the host due to changes in the microbiota and may contribute to the current increased incidence of ailments including inflammatory bowel diseases as well as multiple age-related diseases.

## Introduction

Improper regulation of sugars such as glucose can lead to the development of type 2 diabetes, obesity, inflammatory bowel diseases, cardiovascular disease, and neurodegenerative diseases which it is estimated comprise 10-20% of the worldwide population. These numbers are rising in large part due to the increased amounts of dietary sugar particularly, in the western diet^1^.

Importantly, recently, the human intestinal microbiota has been shown to be affected by diet and influences overall health. In fact, studies suggest that a high-sugar diet can alter the gut microbiota leading to age-associated illness^2,3^. However, mechanistic insight into the critical role of the human intestinal microbiota and the effects of chronic sugar ingestion has not been possible due to the complexity of the human microbiome which contains hundreds of types of organisms. Additionally, much of the data for human studies comes from feces samples which represent only a small portion of the microbes lining the large intestine.

Here, we use an interspecies *Caenorhabditis elegans/Escherichia coli* (*C. elegans*)/(*E. coli*) research platform to simplify direct assessment of importance of the microbiota, and how it is influenced by a high sugar diet. As bacterivores, *C. elegans* have an obligatory symbiotic relationship with microbes as their food source. Interestingly, for normal laboratory growth, *C. elegans* only require one bacterial strain. Therefore, this system allows for univariable analysis of both the intestinal microbiota and the host.

Addition of glucose to the *C. elegans* bacterial diet results in changes in lifespan and healthspan ^4–8^. These studies add glucose either directly to or on top of the agar plate. Therefore, the additional glucose is in contact with both bacteria and *C. elegans* and results in decreased lifespan, reduced healthspan (locomotion), reduced fecundity and changes in fat storage^4–8^.

To separate the effects of glucose on *C. elegans* and bacteria, previously, we developed a new experimental procedure where *E. coli* is incubated with glucose then heat-killed, which results in a glucose fed bacterial diet. Here, we pre-treat the bacteria with glucose but use live bacteria as the food source that becomes the microbiota, and results in a glucose-fed microbiota which we believe mimics the microbiota of a person on a western high added sugar diet.

## Results

Previously ^9^, our data revealed that bacterial processing of added dietary glucose was required for the negative physiological response of the host. To expand these results, we characterized the sensory response of *C. elegans* and performed chemotaxis assays (Fig. 1a). Initially, animals avoided glucose, however, after 24 hours of incubation with *E. coli,* glucose becomes attractive (Fig. 1a). Testing more frequently revealed that the chemotaxis response to glucose is progressive and time-dependent; (Supplemental Fig. 1a). In a similar set-up, where sucrose was added to the bacteria, no significant change was observed at either time point. This is an important control as *E. coli* do not process sucrose ^10^. Therefore, our data illustrate the importance of documenting the time *E. coli* are in contact with glucose prior to addition of *C. elegans*, for studies with an added glucose diet.

**Fig. 1:**
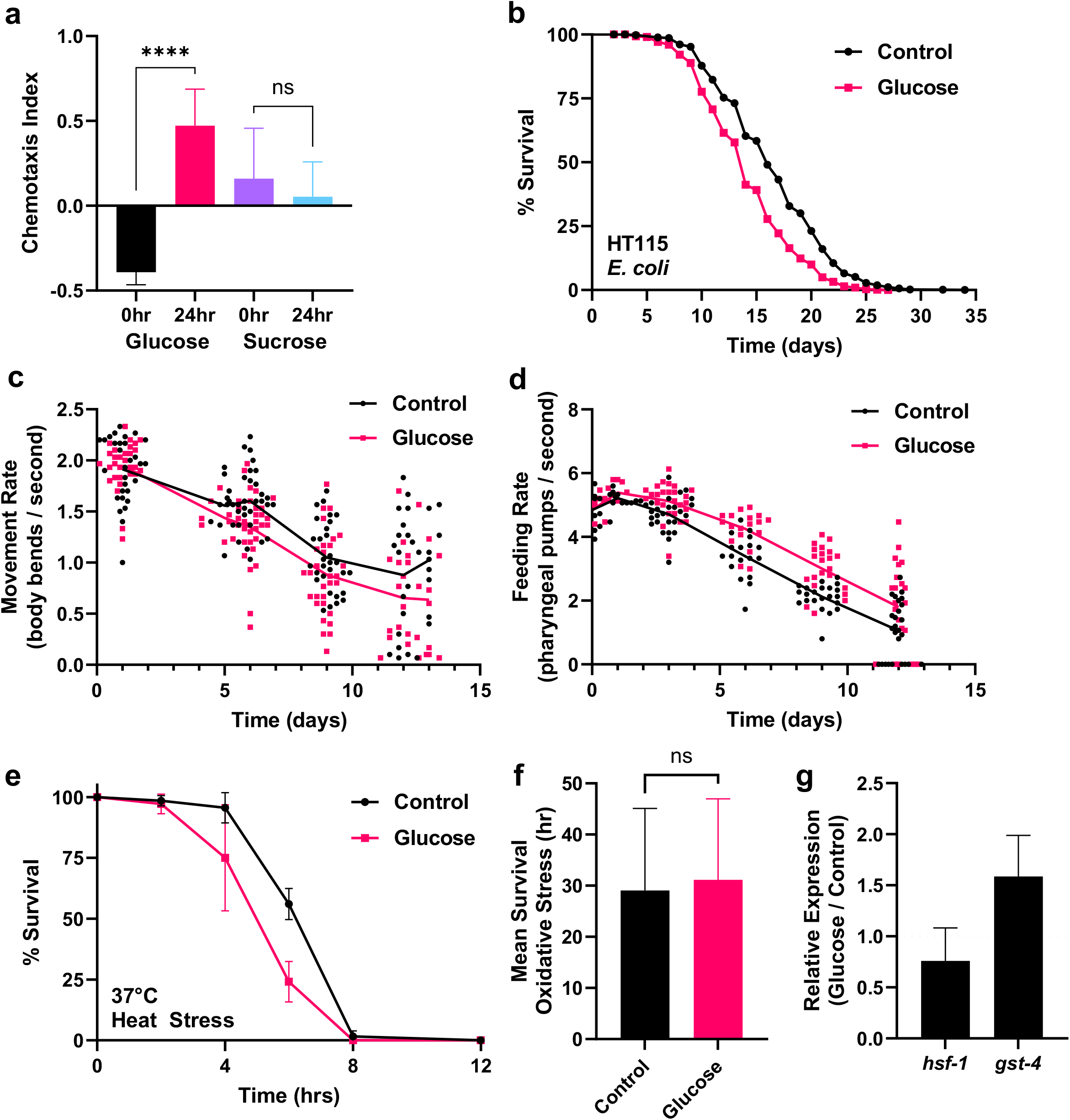
Bacterial incubation with Glucose is required for changes in behavior, lifespan, and health consequences. (a) Chemotaxis Index of *C. elegans* to glucose or sucrose incubated with *E. coli* for 0 or 24 hrs. (b) Lifespan assay of *C. elegans* with glucose fed HT115 *E. coli.* (c) Healthspan-movement in liquid of *C. elegans* growing with a control or glucose fed microbiota over time. (d) Feeding rate/pharyngeal pumping of *C. elegans* growing with a control or glucose fed microbiota. (e) Resistance to heat stress of *C. elegans* grown for 6 days with a control or glucose fed microbiota. (f) Resistance to oxidative stress of *C. elegans* grown for 6 days with a control or glucose fed microbiota. (g) Relative RT-qPCR expression of *hsf-1* and *gst-4* genes in *C. elegans* grown with a control or glucose fed microbiota for 6 days.

Since bacterial processing of glucose affects the attraction of *C. elegans* to *E. coli,* next we measured glucose levels of animals growing with or without bacteria, on agar plates with or without 2% added glucose. Without bacteria, *C. elegans* have a 38% increase in glucose when comparing the 2% glucose and 0% media. In contrast, animals growing on agar plates with *E. coli* have a significant 345% increase in internal glucose when comparing the 2% glucose and 0% media (Supplemental Fig. 1b). These results further indicate the bacteria aids in the animals’ ability to uptake glucose.

Next, to better mimic the mammalian intestine under the duress of a Western high sugar diet, we modified the protocol ^9^ such that *E. coli* are exposed to glucose for three days, then plated to serve as the diet for the host *C. elegans*. The live bacteria will inhabit *C. elegan*s and become the intestinal microbiota resulting in a glucose fed microbiota. Consistently, we saw that the glucose fed microbiota reduced animal lifespan (Fig. 1b). Increased amounts of glucose results in further shortening of lifespan (Supplemental Fig. 1c). From this, we chose 0.8% glucose for the rest of the manuscript.

We tested the effect of glucose on multiple non-pathogenic *E. coli* strains. As shown in Supplemental Figs 2a-c, in additional *E. coli* strains, consumption of live glucose fed OP50, HB101, or BW25113 results in a significant decrease in lifespan. Statistics of the lifespan experiments can be found in Supplementary Table S1. Given that the response to glucose was universal, the remainder of the manuscript used HT115.

To further evaluate how the added glucose affects the aging process, we examined the animals’ healthspan while aging with a glucose fed microbiota. As shown in Fig. 1c, healthspan as measured by movement rate in liquid, also termed body bends/swimming/thrashing, was reduced at all ages after 5 days but not to the level of animals consuming a high glucose diet. These differences may be attributed to the fact that the glucose concentration is a fraction of that seen in other studies. Additionally, after 3 days, there was a significant increase in the feeding rate measured by pharyngeal pumping ^11,12^ (Fig. 1d). The attraction to sugary food is unlikely to be the reason for increased pumping since as our data shows, this is a delayed response. Therefore, examining both healthspan and lifespan showed immediate and long-term consequences of the glucose fed microbiota.

Concurrently, while scoring animal survival on the glucose fed bacteria, avoidance of the bacterial lawn was scored since animals avoid xenobiotic or pathogenic bacteria ^13^. Animals consistently showed less avoidance of the glucose fed bacteria and remained on the bacterial lawn more than the control animals (Supplemental Fig. 2d).

We then tested the ability of animals aging with a glucose fed microbiota to maintain homeostasis. Animals aged 6 days with a glucose fed microbiota show a significant 25% reduction in heat stress resistance (Fig. 1e) but no significant change in oxidative stress resistance (Fig. 1f). Interestingly, RT-qPCR of animals aged for 6 days with the glucose fed microbiota showed reduced expression of a key transcription factor for thermotolerance, heat shock factor-1, *hsf-*1 ^14,15^ and increased expression of *gst-4*, glutathione-S-transferase-4, a marker for oxidative stress, (Fig. 1g). One possibility is that animals with a glucose fed microbiota have reduced heat resistance due to suppression of *hsf-1* gene expression. Accordingly, increased gene expression of *gst-4* would allow for no change in oxidative stress resistance.

Animals with a glucose fed microbiota show a decrease in lifespan as well as an increase in feeding. Previously, a connection between lifespan, food intake and the intestinal microbes was observed such that lifespan is extended on killed bacteria ^16^. Moreover, bacterial overgrowth has been shown to be a cause of death in *C. elegans* ^16^. To confirm that the bacteria are colonizing the intestine, we examined growth of bacteria within the aging *C. elegans* intestine using a fluorescently tagged *E. coli* strain, OP50-GFP. When fed OP50-GFP as *C. elegans* aged, GFP accumulates significantly within the intestine (Supplemental Fig. 3a-b). We then isolated and quantified the bacteria residing within the *C. elegans* intestine measuring the colony forming units (CFU). As a function of age, intestinal bacterial density increases similarly to the fluorescence quantification (Supplemental Fig. 3c).

To ensure that bacterial proliferation was not linked to the GFP tag, we examined animals with a glucose fed microbiota consuming HT115 *E. coli*. As shown in Fig. 2a, intestinal CFU increased at 6 days and significantly increased at 12 days. Therefore, these data show that *C. elegans* exhibit intestinal bacterial packing/overgrowth within the animal as a function of both diet and age. Interestingly, there was no correlation between the mean pumping rate and intestinal bacterial density as the animals aged showing that the increased CFU was not due to increased consumption (Supplemental Fig. 3d).

**Fig. 2:**
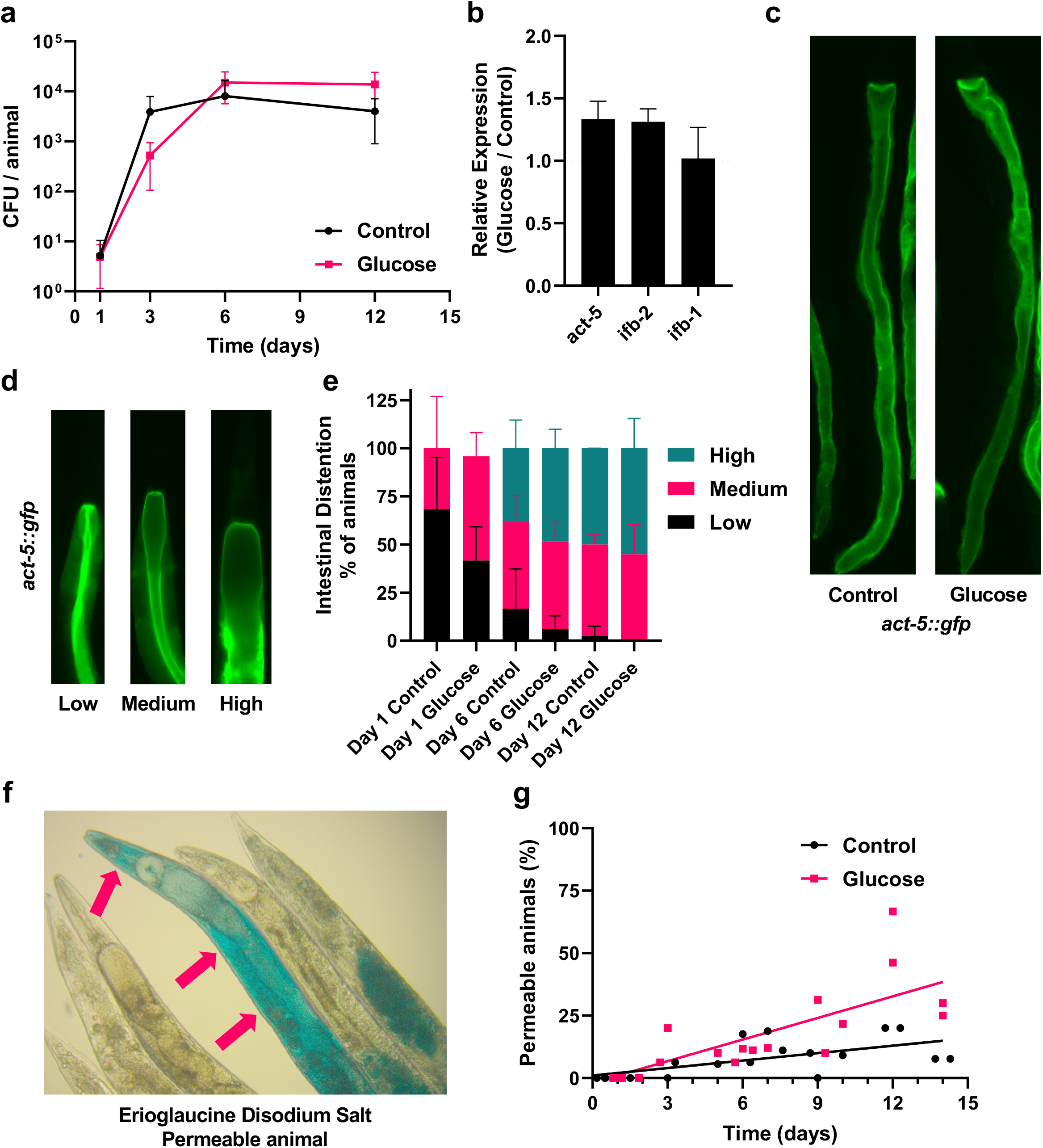
Glucose fed microbiota promotes intestinal bacterial growth, distention, and permeability. (a) Bacterial density of *E. coli* inside *C. elegans* after growing with a control or glucose fed microbiota for 1-12 days. (b) Relative RT-qPCR expression of *act-5, ifb-2, and ifb-1* in *C. elegans* with a control or glucose fed microbiota grown for 6 days. (c) Photographs of *act-5::gfp* growing with control or glucose fed microbiota for 12 days. (d-e) Examples of *act-5::gfp* with low, medium, or high levels of intestinal distention and quantification of *act-5::gfp* intestinal distention after growing with a control or glucose fed microbiota for 1, 6, or 12 days. (f) Example of “Smurf” staining with erioglaucine disodium salt dye within *C. elegans* aged 14 days, arrows indicate the body cavity. (g) Quantification of “Smurf” staining in the body cavity of *C. elegans* growing with a control or glucose fed microbiota for 0-14 days.

Previously, *hsf-1,* has been shown to regulate lifespan and thermotolerance similar to animals with a glucose fed microbiota. *hsf-1* also modifies the actin cytoskeleton ^14,17,18^. Therefore, we next questioned whether the glucose fed microbiota affected the actin cytoskeleton and its function in maintenance of the integrity of the intestinal epithelium. We first examined three genes that function to maintain the intestinal epithelium barrier, the <u>act</u>in *act-5*, and the intermediate <u>f</u>ilament proteins <u>b</u>, *ifb-1*, and *ifb-2.* These three genes colocalize on the apical border of the intestinal epithelium, where *act-5* functions as the primary actin filament within the intestinal microvilli, with *ifb-1* and *ifb-2* as intermediate filament proteins which function to anchor *act-5* on the apical border of the intestinal epithelium ^19^. RT-PCR of day 6 animals with a glucose fed microbiota reveals both *act-5* and *ifb-2* were slightly upregulated (Fig. 2b). In a second series of experiments, we examined *act-5::gfp* transgenic animals aged with and without a glucose fed microbiota (Fig. 2c) and quantification showed no significant whole body fluorescence intensity changes in *act-5::gfp* as the animals aged (Supplemental Fig. 4a). However, as animals aged overall fluorescence steadily declined from day 3 to day 12, indicating a change in the actin cytoskeleton in older animals consistent with previous studies ^20^.

Further analysis of *act-5::gfp* expression in animals with and without a glucose fed microbiota revealed bacterial distention in the aging upper intestine. We classified the upper intestine bacterial distention into three levels: low, medium, and high (Fig. 2d), Quantification of animals as they aged revealed that as a function of age, there was greater bacterial distention. Additionally, our analysis showed day 12 control animals had a similar profile to day 6 animals with a glucose fed microbiota, suggesting premature aging (Fig. 2e). Overall, the proportion of animals scored as medium and high levels of bacterial distention was elevated in the animals with a glucose fed microbiota.

In another series of experiments, we examined the integrity of the intestinal epithelium using the “Smurf” assay, originally used in *Drosophila* and modified successfully for use in *C. elegans* ^21^. The Smurf assay is a staining procedure using erioglaucine disodium salt dye which is taken up by permeable membranes but is excreted by healthy animals, with an example of a positively stained aged animal in Fig. 2f. As expected, young animals showed little permeability in the intestinal barrier, however at 9 days the glucose fed microbiota confers a significant increase in permeability (Fig. 2g). By day 12, ∼56% of animals with the glucose fed microbiota exhibited a Smurf phenotype compared to ∼20% with a control microbiota. Therefore, our data reveal animals with a glucose fed microbiota have increased intestinal epithelial disruption and intestinal bacterial overgrowth.

In humans, illnesses such as small intestinal bacterial overgrowth (SIBO) are characterized by excessive growth of bacteria in the small intestine ^22^. Interestingly, multiple homeostatic systems control the bacterial population in the small intestine but conditions such as aging and diabetes disrupt these mechanisms and lead to SIBO ^22^. Bacterial over proliferation illnesses such as SIBO leads to inflammation and activation of the immune response. Therefore, we next examined if there was any connection between bacterial overgrowth caused by the glucose fed microbiota, inflammation, and the immune response.

In *C. elegans*, there is no true inflammation. However, it is well documented that similar to mammalian phagocytes, *C. elegans* produce reactive oxygen species (ROS) in response to infection ^23^. We measured whole animal ROS and found ROS was increased with a glucose fed microbiota at both days 1 and 6 (Fig. 3a). We questioned if the expression of *gst-4::gfp* was also regulated by the bacteria and so we heat killed the glucose fed bacteria and found that animals are not able to elicit the same upregulation of *gst-4* without live bacteria (Fig. 3b-c). Therefore, animals with a live glucose fed microbiota have an increase in oxidative stress reported by *gst-4::gfp* as well as increased ROS.

**Fig. 3:**
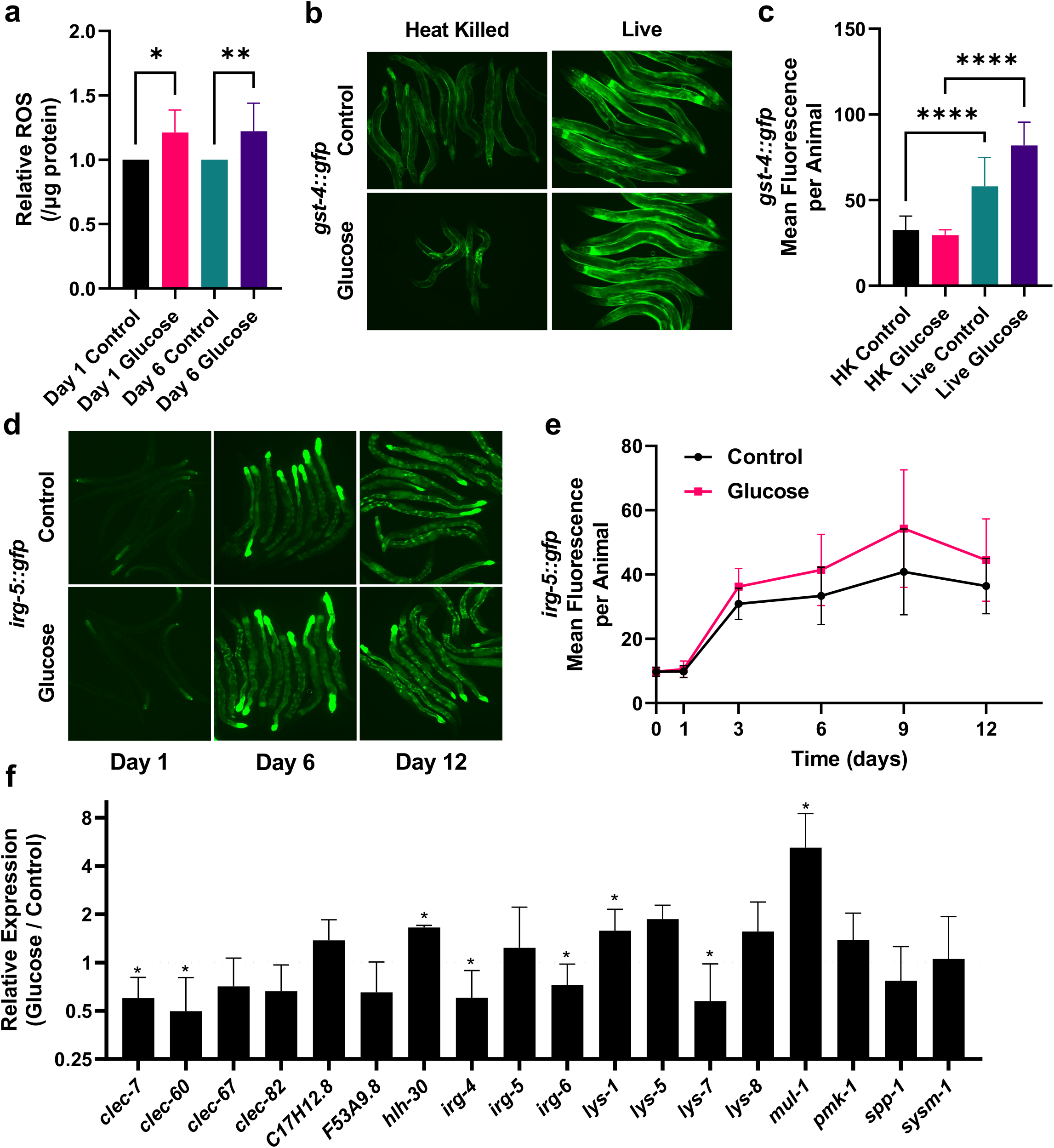
A glucose fed microbiota promotes oxidative stress. (a) Reactive Oxygen Species (ROS) measurement of *C. elegans* growing with a control or glucose fed microbiota for 1 or 6 days. (b) Photographs of *gst-4::gfp* animals after growing with either a heat killed, or live control and glucose fed microbiota for 6 days. (c) Quantification of *gst-4::gfp* whole body fluorescence after growing with either a heat killed (HK) or live control and glucose fed microbiota for 6 days. (d) Photographs of *irg-5::gfp* after growing with a control or glucose fed microbiota for 1, 6, or 12 days. (e) Quantification of *irg-5::gfp* whole body fluorescence after growing with a control or glucose fed microbiota for 1-12 days. (f) Relative RT-qPCR expression of a panel of innate immune reactive genes in *C. elegans* grown with a control or glucose fed microbiota for 6 days.

**Fig. 5:**
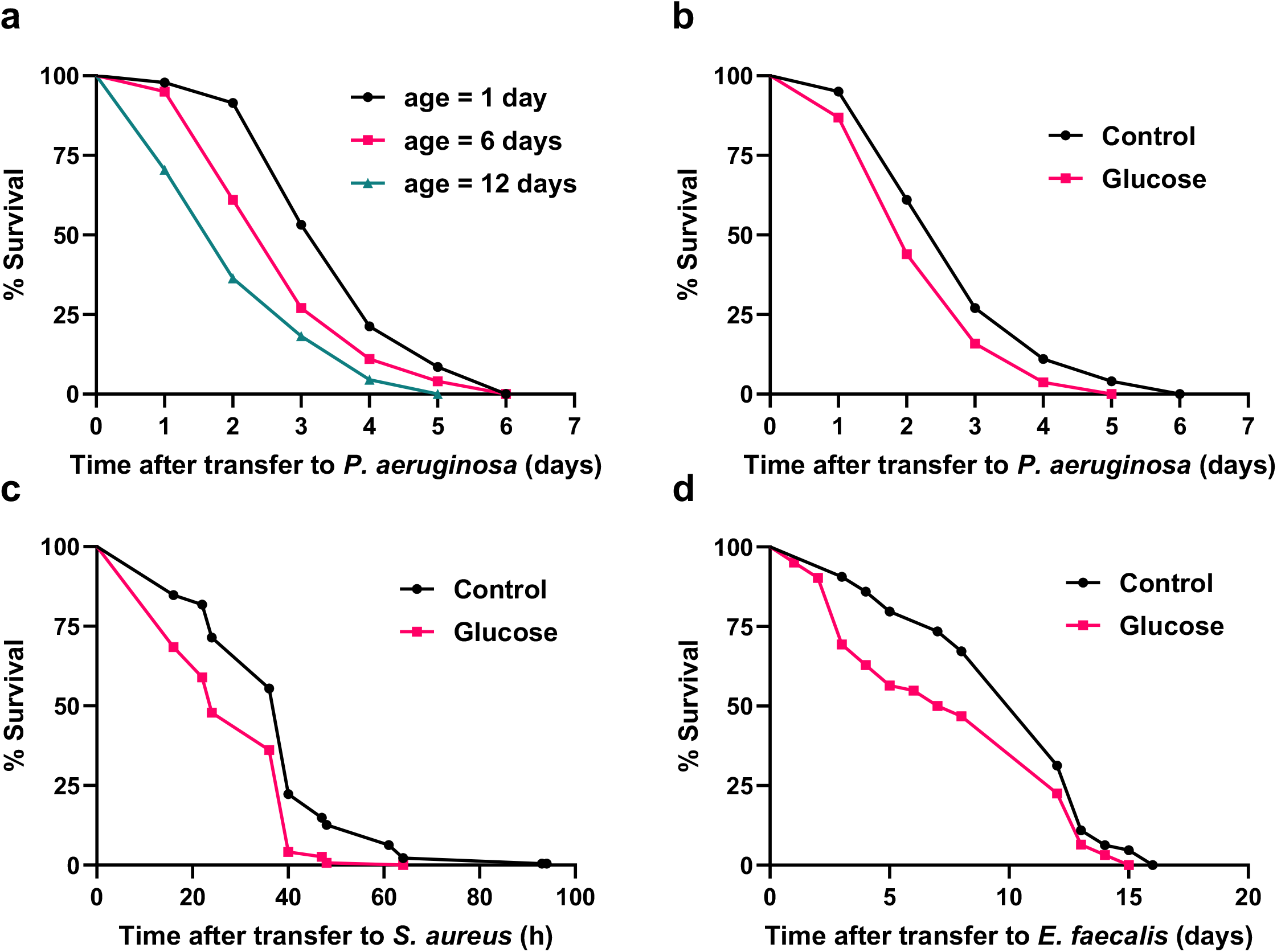
Aging with a glucose fed microbiota results in susceptibility to multiple bacterial pathogens. (a) Survival of *C. elegans* aged 1, 6, or 12 days with a control or glucose fed microbiota before transfer to *P. aeruginosa*. (b) Survival of *C. elegans* on *P. aeruginosa* after growing with a control or glucose fed microbiota for 6 days. (c) Survival of *C. elegans* on *S. aureus* after growing with a control or glucose fed microbiota for 6 days. (d) Survival of *C. elegans* on *E. faecalis* after growing with a control or glucose fed microbiota for 6 days.

The bacterial overgrowth and increased ROS production when animals have a glucose fed microbiota may indicate that the animals are perceiving a pathogen, which in *C. elegans* is sensed by the innate immune system. When *C. elegans* are exposed to a bacterial pathogen including *Pseudomonas aeruginosa* (*P. aeruginosa*)*, Staphylococcus aureus*(*S. aureus*), *Yersinia pestis* and *Enterococcus faecalis* (*E. faecalis*), gene expression of immune effectors including CUB-like genes, lectins, lysozymes, PUFA genes more than double ^24–27^. To examine the relationship between innate immunity and the glucose fed microbiota, we first examined a fluorescent transcriptional reporter for an immune response gene, *irg-5* ^28,29^ since expression of *irg-5* is dependent on multiple immune pathways ^27^. After 3 days, the glucose fed microbiota increased expression of *irg-5::gfp* (Fig. 3d-e). We then examined innate immune effector genes previously linked to a bacterial pathogenic response including C-type lectins, lysozymes, mucin, MAP kinase, and other infection related/innate immune genes ^24–26^. As shown in Fig. 3f, gene expression data showed only ∼50% of the innate immune genes tested were significantly changed. Furthermore, some genes had reduced expression with a glucose fed microbiota, unlike animals exposed to bacterial pathogens such as *Pseudomonas aeruginosa* (*P. aeruginosa*) or *Enterococcus faecalis* (*E. faecalis*) ^24–26,30^. Therefore, animals with a glucose fed microbiota show signs of a bacterial infection (increased CFU/bacterial overgrowth, increased ROS production and increased oxidative stress) ^24–27^, However, we suggest that rather than a bacterial pathogen infection, this is dysregulation of immune function.

To further examine the immune dysregulation in combination with increased intestinal epithelial permeability, next, we tested resistance to multiple bacterial pathogens of animals aging with a glucose fed microbiota. Animals aged for 1, 6, or 12 days on a control diet before exposure to the bacterial pathogen *P. aeruginosa* showed an age dependent decrease in survival (Fig. 4a). Then, animals aged 6 days with a glucose fed microbiota showed a significant reduction in survival when exposed to the bacterial pathogen *P. aeruginosa* (Fig. 4b). Similarly, animals with a glucose fed microbiota showed a significant reduction in survival when exposed to the bacterial pathogen, *S. aureus* (Fig. 4c). Animals with a glucose fed microbiota also showed markedly reduced resistance to a third bacterial pathogen, *E. faecalis* (Fig. 4d). Therefore, as animals age, the ability to respond to bacterial pathogens declines, and across multiple bacterial pathogens, animals with a glucose fed microbiota show vulnerability and decreased resistance to bacterial pathogenic infection.

In summary, our results reveal that animals with a glucose fed microbiota show lifelong significant alterations in both lifespan and healthspan. Our data revealed that the health consequences of a glucose fed microbiota occur progressively. Initially, animals show an increase in feeding, mild gut distention, and increased ROS. By day 3, animals show immune dysregulation and reduced movement. On day 6, there is increased bacterial accumulation, increased intestinal distention, increased ROS, and increased bacterial pathogen susceptibility coupled with reduced ability to survive stress, and immune dysregulation. By day 9, there is significant increased permeability of the intestinal epithelium. By day 12, the animals with a glucose fed microbiota show a significantly increased chance of death. Interestingly, all phenotypes associated with a glucose fed microbiota are age accelerated, occurring days before the mean survival time. This suggests that alleviating these symptoms could change the course of an animal’s health and survival.

Overall, our data reveal that two important factors, diet and the aging process contribute to affect lifespan and healthspan. We clearly show that as animals age with a glucose fed microbiota, they have reduced lifespan and become unhealthy as exemplified by an increase susceptibility to multiple bacterial pathogens. In a society where food intake exceeds the necessary nutritional requirements of an individual and surpasses the rate limiting import of food, ingested nutrition by a host (human or *C. elegans*) is inadvertently exposed to and metabolized by intestinal microbiota. We show the critical importance of a healthy microbiota throughout the aging process. Together, our data show the potential link between diet and bacterial infection and possibly represent a new avenue for therapeutics.

## Acknowledgement

We are grateful to members of the Tissenbaum and Finkel lab for advice and suggestions, Susan Lee and Evelyn Caez for technical support, Dr. David Greenstein for advice and support, Emily Troemel for providing strains and Read Pukkila-Worley for providing strains, providing primer sequences and help with the *P. aeruginosa* experiments. Some of the *C. elegans* strains were kindly provided by the *Caenorhabditis* Genetics Center, which is funded by NIH Office of Research Infrastructure Programs (P40 OD010440). H.A.T. is a William Randolph Hearst Investigator. This project was funded by a grant from the NIH/NIA (5R21AG067317) to H.A.T. S.F. is supported (by a grant from the U.S. Army Research Office (W911NF1210321) and H.A.T. is supported in part by an endowment from the William Randolph Hearst Foundation.

## Methods

### Strain maintenance

All *C. elegans* strains were maintained at 20°C using standard *C. elegans* techniques^31^. Wild type N2 *C. elegans* was used for most experiments. The following transgenes were used: AY101 *acIs101* [F35E12.5p::GFP + rol-6(su1006)], ERT60 *jyIs13* [*act-5p*::GFP::ACT-5 + *rol-6(su1006*)] II., CL2166 *dvIs19* [(pAF15)*gst-4p*::GFP::NLS] III.

### Chemotaxis Assay

The chemotaxis assay was adapted from ^32,33^. Standard NGM 60mm plates were divided into 4 quadrants, with a 10mm diameter circle in the center, and 4 dots (one in each quadrant 1mm from the edge of the plate). 10μL of 2M D-Glucose (G8270-1KG, Sigma Aldrich) or 2M Sucrose (S0389-500G, Sigma Aldrich) solubilized in M9 was pipetted onto the dots of two opposing quadrants with control (M9) on the last two quadrants and allowed to dry. Then, 10 μL of OP50 *E. coli* was spotted on top of the dried treatment and plates were either left to incubate at room temperature for 24 hours or used immediately after bacteria had dried. Well-fed *C. elegans* adults (approximately 25-50/plate) were picked from stock OP50 plates, washed three times with M9, and picked onto the center of all assay plates. After one hour, plates were placed at 4°C for 15 minutes to slow/immobilize animals for ease of scoring and the location of the animals recorded. The chemotaxis index refers to the (# of animals on treatment - # of animals on control) / total # of animals. Experiments were performed on 3 biological replicates in triplicate, and statistics calculated using an unpaired t-test with Graphpad Prism 10 (Graphpad Software).

### Measurement of Total Glucose

Total glucose of *C. elegans* lysates was measured using a Glucose Assay Kit (GAGO20-1KT, Sigma-Aldrich) according to the manufacturers’ protocol. NGM plates with either 0% or 2% added glucose were seeded with/without 250uL of OP50 *E. coli* and then dried for 2 days. Approximately 100 *C. elegans* L4s were picked onto each plate and grown at 20°C for 2 days. Animals were then picked off plates, washed 2x with M9 buffer, and then frozen at −80°C. Frozen pellets were reconstituted in 7 volumes of RIPA (150mM NaCl, 50mM Tris pH 7.4, 1% Triton X-100, 0.1% Sodium Dodecyl Sulfate, 1% Sodium Deoxycholate) buffer freshly mixed with Protease Inhibitor Cocktail (P2714-1BTL, Sigma Aldrich). Samples were then lysed on ice for 30 seconds total using a Microson XL2000 probe sonicator at 25% power. Each sample was also analyzed with a Pierce Coomassie Plus (Bradford) assay (23236, Thermo Scientific) and Glucose values were normalized by protein values. Experiments were performed on 3 biological replicates in duplicate, and statistics were calculated using an unpaired t-test using Graphpad Prism 10 (Graphpad Software).

### Lifespan Assay

*C. elegans* L4s were picked onto control or glucose fed *E. coli* plates and grown at 20°C (∼25 animals per 60mm plate). Animals were transferred to new plates every 2 days and scored by gently tapping with a platinum wire pick daily while producing progeny and then every 2-3 days. Animals that did not respond were scored as dead. Animals that were lost, desiccated, bagged, or died from vulva bursting were censored from the analysis. Each Figure shows cumulative data collected from at least 2 biological replicates in triplicate. Statistics were calculated by a two-tailed unpaired t-test using Graphpad Prism 10 (Graphpad Software) and shown in Supplementary Table 1.

### Glucose-fed Microbiota

Stabs of stock frozen cultures of OP50, HT115(DE3)-L4440 (referred to as HT115), HB101, and BW25113 strain *E. coli* were used to inoculate 50mL LB cultures to start each culture. HT115 was grown in the presence of ampicillin, HB101 in the presence of streptomycin, and OP50 and BW25113 with no antibiotics. Bacterial cultures were grown in LB with either sterile filtered ddH_2_O (control) or 0.8% D-Glucose aerobically at 37°C on a shaker plate for 3 days, then pipetted onto NGM plates and allowed to dry for 2 days. After drying, *C. elegans* were picked onto the plates and aged for analysis.

### Movement in liquid

*C. elegans* L4s were picked onto control or glucose fed *E. coli* plates and grown at 20°C for 1, 5, 6, 9, 12, or 13 days. Individual animals were picked onto an unseeded NGM plate, then 10μL of M9 buffer was pipetted onto each animal, and the number of body bends was counted over 30 seconds. Experiments were performed on 3 biological replicates with 10 animals each, Statistics were calculated by an unpaired t-test using Graphpad Prism 10 (Graphpad Software).

### Feeding Rate

*C. elegans* L4s were picked onto control or glucose fed *E. coli* plates and grown at 20°C for 0, 1, 3, 6, 9, and 12 days and counted manually for pharyngeal pumping contractions over 15 seconds. Experiments were performed on 3 biological replicates with 8 animals each. Statistics were calculated by an unpaired t-test using Graphpad Prism 10 (Graphpad Software).

### Resistance to Oxidative Stress

*C. elegans* L4s were picked onto control or glucose fed *E. coli* plates and grown at 20°C for 6 days. Then, 1mL of 250mM Paraquat (Methyl viologen dichloride hydrate, 856177-1G, Sigma Aldrich) solution was added to the 60 mm NGM plates and plates were put on a shaker for 1 hour, followed by 1.5 hours in a laminar flow hood to ensure plates were dry and the paraquat evenly distributed. Animals were then transferred to paraquat plates at 20°C and scored twice daily for survival. Animals were touched with a platinum wire, and those that did not respond were scored as dead. Experiments were performed on 5 biological replicates with 25-30 animals each. Statistics were calculated by an unpaired t-test using Graphpad Prism 10 (Graphpad Software).

### Resistance to Heat Stress

*C. elegans* L4s were picked onto control or glucose fed *E. coli* plates and grown at 20°C for 6 days, then transferred to 37°C for either 2, 4, 6, 8, 12, or 16 hours and scored for survival. Animals were touched with a platinum wire, and those that did not respond were scored as dead. Percent survival was calculated for each time point and animals were not placed back to 37°C after scoring. Experiments were performed on 3 biological replicates with 30-35 animals each, and statistics were calculated by an unpaired t-test using Graphpad Prism 10 (Graphpad Software).

### Bacterial Density (CFU)

*C. elegans* intestinal bacterial density measurement was done following ^34^. *C. elegans* L4s were picked onto control or glucose fed *E. coli* plates and grown at 20°C for 1, 3, 6, or 12 days. Animals were counted and picked off plates, washed 3x with M9 buffer and then immobilized 0.25mM levamisole and immersed into a 3% bleach solution to sterilize the exterior of the animal for 5 minutes. Animals were then washed 3x with M9 buffer, resuspended in 1% Triton X-100, then ground with a pellet pestle. The samples were centrifuged at 14,000 x g for 10 min at 4°C then had supernatant removed, and the pellet was resuspended in 500uL M9 buffer. A series of 10-fold dilutions were then made using ddH_2_O from 10^−1^ to 10^−5^ dilution. In a laminar flow hood, for each sample in triplicate, 50uL was pipetted onto a 35mM LB agar plate and spread across the surface. After plates dried, they were moved to 37°C for 16 hours, followed by colony counting. Only plates with colony counts between 30-300 were included, then they were multiplied by the dilution factor and divided by number of animals used to produce CFU/animal. Experiments were performed on 3 biological replicates with 45-55 animals each, and an unpaired t-test was done using Graphpad Prism 10 (Graphpad Software).

### ROS Quantification

Measurement of *C. elegans* Reactive Oxygen Species (ROS) was done following ^35^. *C. elegans* were picked onto control or glucose fed *E. coli* plates and grown at 20°C for either 1 or 6 days. Then, 100 animals were picked off plates and washed 3x with M9 buffer. At the last wash, the animal mix was left in 100uL which was evenly distributed into two wells of a 96-well plate. Either 50uL M9 buffer (control) or 50uL of 50uM H_2_DCFDA was added to the well, then the plate was wrapped with aluminum foil and placed on a shaker for 30 minutes. A fluorometer was then used to measure fluorescence at 490 nm and 520 nm. Fluorescence of the blank control was subtracted from all samples and the relative difference in fluorescence calculated. Experiments were performed on 3 biological replicates in duplicate, and statistics were calculated by an unpaired t-test using Graphpad Prism 10 (Graphpad Software).

### Fluorescent Imaging and Quantification

Transgenic *C. elegans* L4s were picked onto control or glucose fed *E. coli* plates and grown at 20°C for 1, 3, 6, 9, or 12 days. Then, animals were mounted onto slides and were imaged using a Hamamatsu ORCA ER camera mounted onto a Zeiss Axioskop 2 plus equipped with a FITC filter. Photographs were exported as .tiff files and quantified using ImageJ 1.52a (https://imagej.nih.gov/ij/index.html) by closely outlining whole individual animals and measuring pixel intensity. Images were captured in monochrome and artificially recolored as green using ImageJ. Experiments were performed on 2-6 biological replicates with 5-10 animals per treatment, and an unpaired t-test using Graphpad Prism 10 (Graphpad Software) was done.

### RNA extraction and RT-qPCR

*C. elegans* L4s were picked onto control or glucose fed *E. coli* plates and grown at 20°C for 6 days. Then, 100 animals were washed off plates with M9 buffer and rinsed twice with ddH_2_O. Total RNA was isolated using TRIzol Reagent with the Direct-zol RNA MiniPrep (Zymo Research). Quality control of the RNA was performed using a Nanodrop (Thermo Scientific) and samples with both an A260/A280 ratio > 2.0 and an A260/A230 ratio > 1.8 were used. First-strand cDNA synthesis was performed on 1.0μg of total RNA using dNTPs, Oligo(dT)_12–18_ and SuperScript III Reverse Transcriptase (Invitrogen). Reverse Transcriptase Quantitative PCR was done using the QuantStudio™ 3 System (Applied Biosystems) with Power SYBR Green PCR Master Mix (Applied Biosystems) per the manufacturer’s instructions, in triplicate. The endogenous control for relative expression normalization was *act-1* for Figs. 1g and 2b, and then *snb-1* for innate immune response genes in Fig. 3f. Sequences of primers can be found in Supplementary Table S2. Relative fold changes of gene expression were calculated using Comparative C_T_ (ΔΔC_T_). Experiments were performed on 3 biological replicates in triplicate, and an unpaired t-test using Graphpad Prism 10 (Graphpad Software) was used.

### Intestinal Distention

ERT60 jyIs13 [*act-5p::GFP::ACT-5 + rol-6(su1006)*] L4s were picked onto control or glucose fed *E. coli* plates and grown at 20°C for 1, 6, or 12 days. Then, animals were mounted on slides and observed on a Zeiss Axioskop 2 plus for bacterial distention of the upper intestine. Distention was classified into three different levels: low, medium, and high. Experiments were performed on 3-6 biological replicates with 5-10 animals per treatment, and an unpaired t-test using Graphpad Prism 10 (Graphpad Software) was done.

### Dye Uptake “Smurf” assay

Assay was adapted from ^21^. *C. elegans* L4s were aged on control or glucose fed *E. coli* for 0, 1, 3, 5, 6, 7, 9, 10, 12, or 14 days at 20°C. Then, animals were picked off plates into an Eppendorf vial with 5% Erioglaucine disodium salt solution added and incubated at RT on a shaker plate for 3 hours. Vials were centrifuged gently, and supernatant removed to 50uL. Using a glass pipet, animals were dispensed onto an OP50 NGM plate for 30 minutes. Animals that did not move or remained in the dye were omitted from further analysis. Then, animals were mounted onto slides and were imaged using a Zeiss AxioCam ERc 5s mounted on a Zeiss Stemi SV11 Apo using bright field light. Experiments were performed on 2-3 biological replicates with 10-25 animals per treatment, and an unpaired t-test using Graphpad Prism 10 (Graphpad Software) was used.

### Pathogen Survival assay

*C. elegans* L4s were picked onto control or glucose fed *E. coli* plates and aged for 1, 6, or 12 days at 20°C. Animals were then picked onto either *P. aeruginosa*, *S. aureus*, or *E. faecalis* plates and scored 1-2 times per day for survival. Animals were scored by gently tapping with a platinum wire pick, those that did not respond were scored as dead. Animals that were lost, bagged, or died from vulva bursting were censored from the analysis. Experiments were performed on 2 biological replicates in triplicate, and statistics were calculated by a two-tailed unpaired t-test using Graphpad Prism 10 (Graphpad Software). Statistics of each experiment are located in Supplementary Table 1.

*P. aeruginosa* assays were performed in the laboratory of Dr Reed Pukkila-Worley. Briefly, plates were prepared using the protocol in ^26,36^. A single colony of *P. aeruginosa* PA14 was inoculated into 3-mL of Luria-Bertani (LB) media and incubated at 37°C for 14 to 15 hours. 10 mL of this culture was added to 35-mm tissue culture plates containing 4 mL of slow kill agar. Plates were incubated for 24 hours at 37°C and 24 hours at 25°C and 0.1 mg/mL 5-fluorodeoxyuridine (FUDR) was added to the media 1 to 2 hours prior to the start of the assay to prevent progeny from hatching. The *P. aeruginosa* survival assays were conducted at 25°C.

*S. aureus* assays were performed in the laboratory of Dr. Javier Irazoqui. *S. aureus* plates were prepared using the protocol described in ^37^. *S. aureus* SH1000 was grown overnight in TSB containing 50 mg/ml Kanamycin. Then 500–1000 ml of overnight culture was uniformly spread on the entire surface of freshly prepared 100 mm TSA plates supplemented with 10 mg/ml Kanamycin. The plates were incubated for 6 hr at 37°C, then stored overnight at 4°C. The plates were warmed to room temperature and *S. aureus* survival assays were conducted at 20°C. Scoring of the added glucose was done blind.

*E. faecalis* plates were prepared using similar methods as standard stock OP50 plates. Stock *E. faecalis* isolated from OMC47 human donors was inoculated into 20mL LB and cultured for 16 hours at 37°C in an aerobic environment. Approximately 300mL of the culture was aliquoted onto standard NGM plates, and then the plates were placed into an anaerobic chamber for 16 hours at 37°C. Plates were then kept at 20°C and *C. elegans* were added.

## Results and Discussion

Since *C. elegans* are bacterivores, they have an obligatory symbiotic relationship with microbes as their food source. We have used this to our advantage in our *C. elegans/E. coli* paradigm. We have previously focused on separating the effects of glucose on *C. elegans* and *E. coli*. To do so, we developed an experimental procedure where the *E. coli* is grown with glucose for 3 days, then heat-killed and used as food resulting in a diet consisting of glucose fed bacteria. In contrast, in this manuscript, we grow the *E. coli* with glucose for 3 days and then use this live *E. coli* as the bacterial diet which becomes the microbiota, and results in a glucose-fed microbiota.

Therefore, the bacteria that become the microbiota have already processed the additional glucose allowing interrogation of the downstream effects.

### Dye Retention in Smurf staining

Further examination of the erioglaucine disodium salt-stained animals revealed that there were dye retention changes in different tissues (pharynx, upper intestine, mid-intestine/vulva, lower intestine) both as animals aged and with a glucose fed microbiota. Although there were relatively little changes in the pharynx or upper intestine, the glucose fed microbiota further increased dye retention within the mid intestine, vulva, and lower intestine over time (Supplemental Figures 4a-e). Dye retention may reveal where chemicals and possibly bacteria may find shelter from excretion. In general, the observed dye retention was increased with age in all tissues, and early age animals show little to no differences with a glucose fed microbiota, however as those animals age greater dye presence was found within the vulva and middle to lower intestines. Dye retention suggests the sites of permeability or where chemicals and possibly bacteria may find shelter from excretion. In general, the observed dye retention was increased with age in all tissues, and early age animals show little to no differences with a glucose fed microbiota.

However, as animals age with a glucose fed microbiota, greater dye retention was seen within the vulva and middle to lower intestines. We suggest examining dye retention as a function of diet and age.

### Nutritional status of animals with a glucose fed microbiota

The nutritional status of animals aged with a glucose fed microbiota is unknown. As shown in Figure 1d, animals with a glucose fed microbiota have an increase in feeding. According to Avery and Shtonda (2003)^1^, an increased feeding rate indicates poor nutritional status. Therefore, animals with a glucose fed microbiota maybe slightly starved. However, our data reveals that animals with a glucose fed microbiota have a shortened lifespan (Figure 1b) and *C. elegans* undergoing dietary restriction have an increase in lifespan^2,3^. This suggests that we are not seeing nutrient restriction but perhaps nutrient malabsorption.

### Nutrient Absorption affected by a glucose fed microbiota

Intestinal microvilli concentration as well as dietary uptake into epithelial tissue within *C. elegans* is determined by *act-5* expression ^4^. Animals with a glucose fed microbiota have increased expression of *act-5*, in addition to the anchoring terminal web intermediate filament *ifb-2*, perhaps suggesting increased intestinal absorption and an adaptation to a lower quality diet (Fig. 3).

Although *act-5::gfp* animals did not show significant differences in total fluorescence with a glucose fed microbiota, animals did show increased intestinal distention as they aged. In addition, with age, *act-5::gfp* animals with a glucose fed microbiota, showed changes in the intensity of fluorescence between the upper and lower intestine. Therefore, together our data reveal the importance of maintaining the integrity of the intestinal epithelium and the influence of age and diet.

---

**Supplementary Figure 1:**
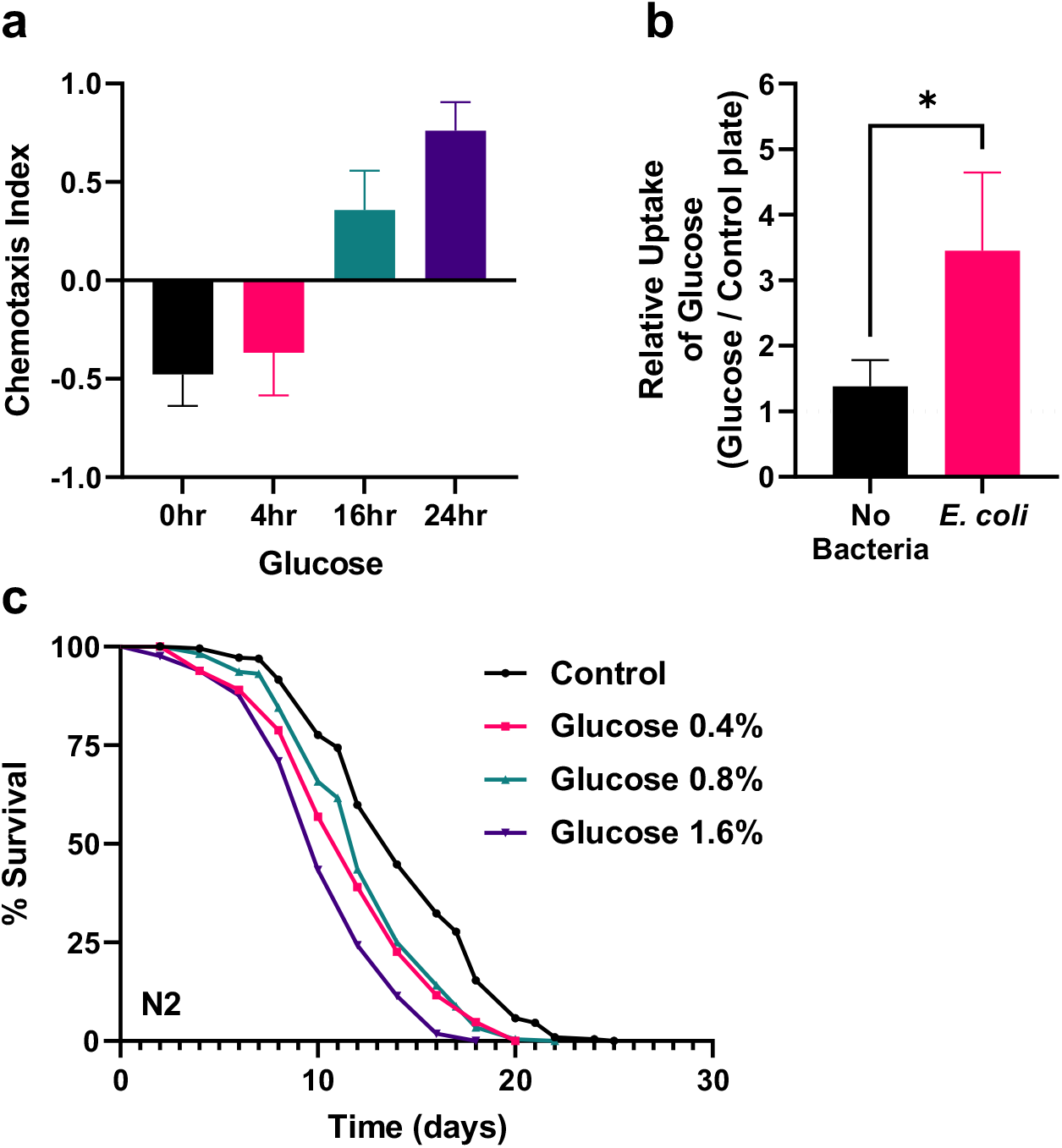
Incubation time and concentration of glucose modulate behavior, glucose uptake, and lifespan. (a) Chemotaxis Index of *C. elegans* showing preference to *E. coli* incubated with glucose for 0, 4, 16, or 24hrs. (b) Whole animal glucose uptake assay of *C. elegans* after growing for 2 days on 2% glucose plates without bacteria or seeded with OP50 *E. coli* compared to respective 0% glucose controls. (c) Lifespan assay of *C. elegans* with 0, 0.4, 0.8, 1.6% glucose fed HT115 *E. coli*.

**Supplementary Figure 2:**
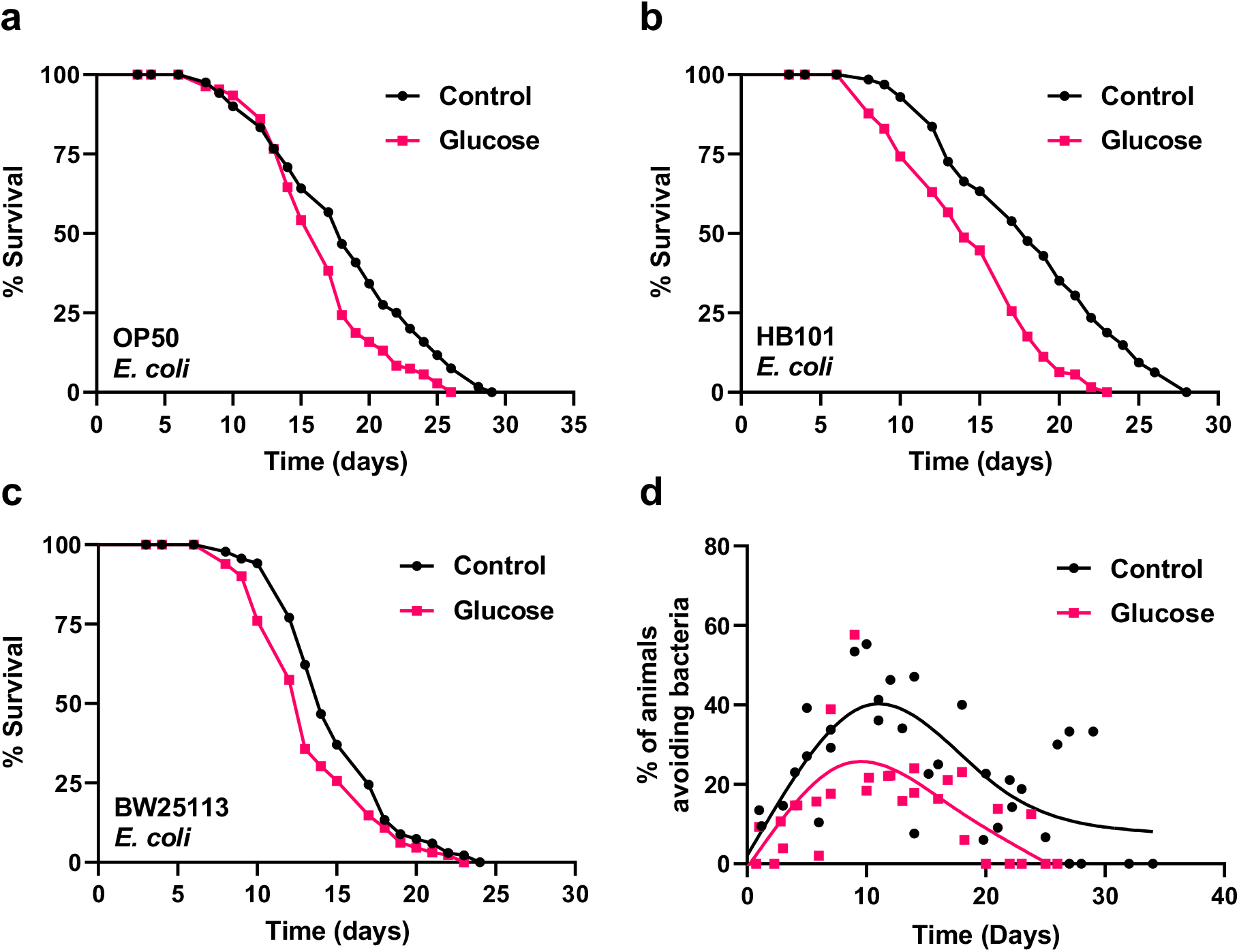
(a-c) Lifespan effect of glucose across multiple strains of *E. coli.* (a) Lifespan assay of *C. elegans* with glucose fed OP50 *E. coli*. (b) Lifespan assay of *C. elegans* with glucose fed HB101 *E. coli*. (c) Lifespan assay of *C. elegans* with glucose fed BW25113 *E. coli*. (d) Avoidance behavior of *C. elegans* on glucose fed HT115 *E. coli* plates throughout course of lifespan experiments, measured by % of animals off bacterial lawn out of total animals per plate, each dot represents a batch of ≤ 30 animals.

**Supplementary Figure 3:**
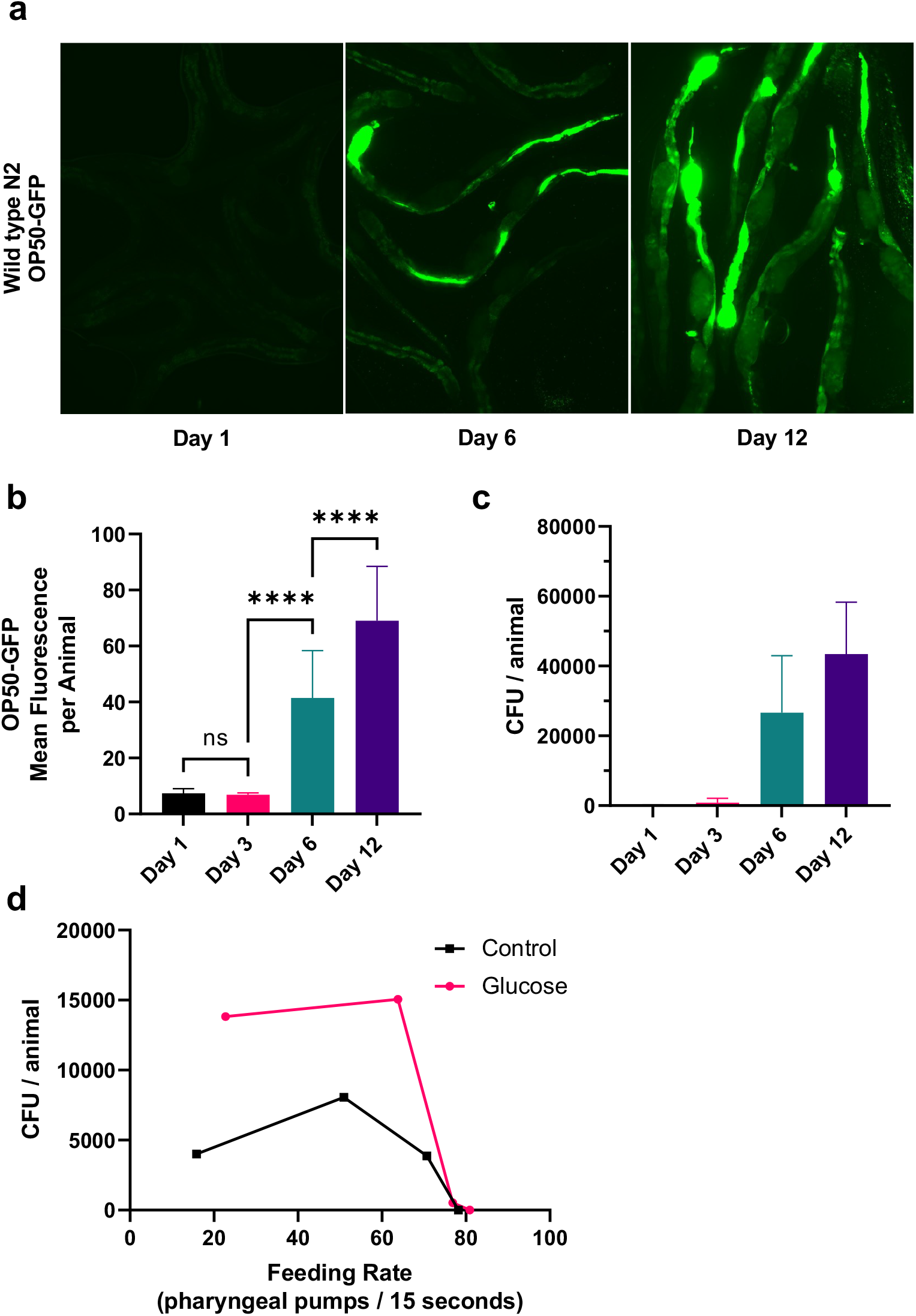
(a-c) Growth of bacteria within aging *C. elegans*. (a) Representative images of OP50-GFP *E. coli* within the intestine of *C. elegans* after feeding for 1, 6, or 12 days. (b) Quantification of OP50-GFP *E. coli* fluorescence within the intestine after feeding for 1, 3, 6, or 12 days. (c) Bacterial density of OP50-GFP *E. coli* isolated from *C. elegans* after feeding for 1, 3, 6, or 12 days. (d) No correlation between the mean pumping rate and intestinal bacterial density as the animals aged with a control or glucose fed microbiota for 1, 3, 6, or 12 days.

**Supplementary Figure 4:**
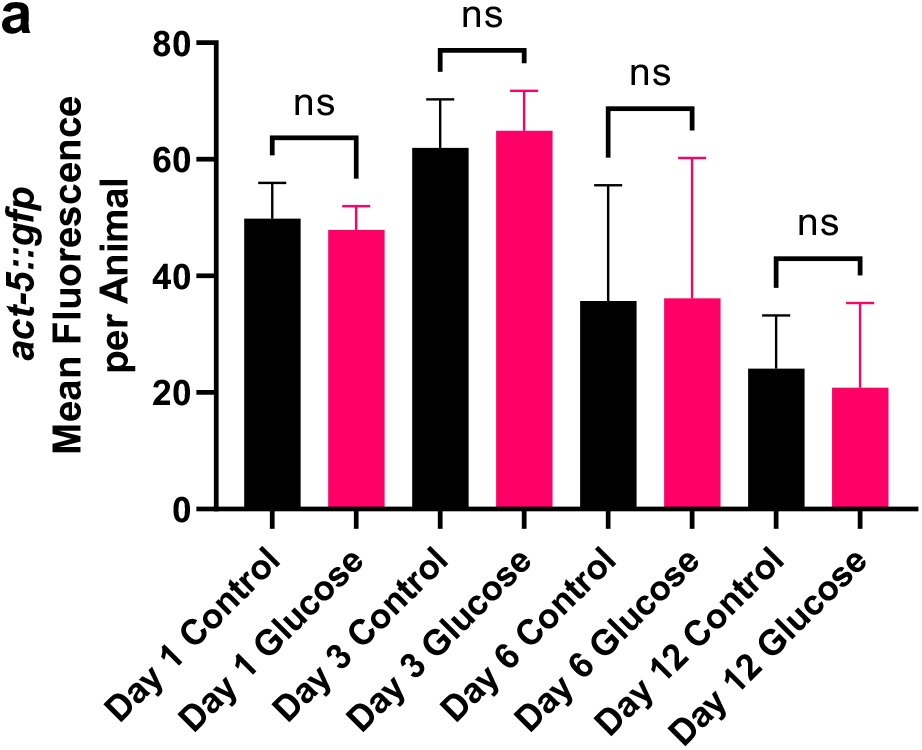
A glucose fed microbiota results in changes in the intestinal epithelial structure. Quantification of *act-5::gfp* whole body fluorescence after growing with a control or glucose fed microbiota for 1, 3, 6, or 12 days.

**Supplementary Figure 5:**
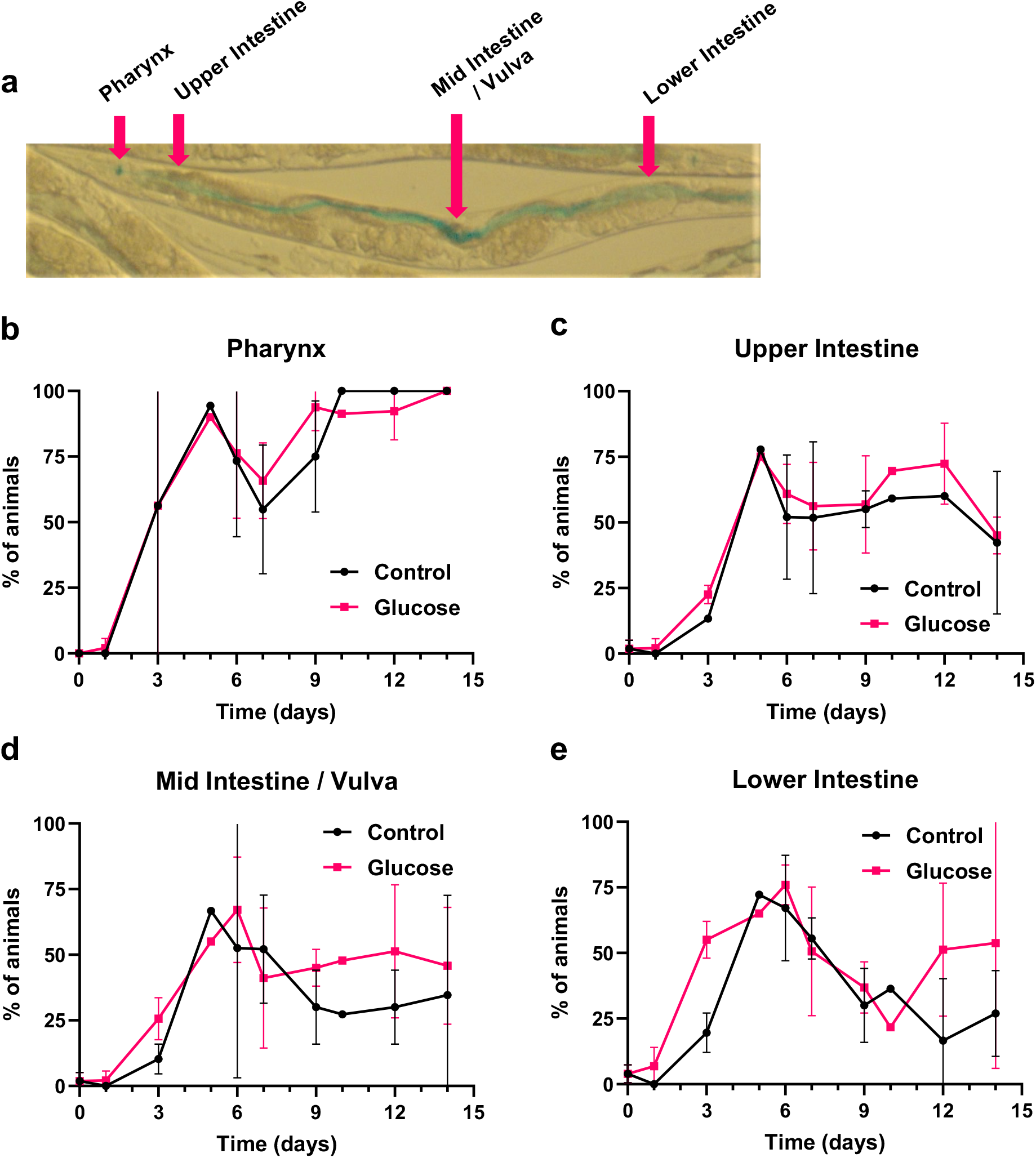
A glucose fed microbiota results in intestinal epithelial permeability. (a) Representative image of *C. elegans* stained with erioglaucine disodium salt dye retention in the pharynx, upper intestine, mid intestine/vulva, and lower intestine. (b) Quantification of erioglaucine disodium salt dye retention in the pharynx of *C. elegans*, each dot represents 10-25 animals. (c) Quantification of erioglaucine disodium salt dye retention in the upper intestine of *C. elegans*, each dot represents 10-25 animals. (d) Quantification of erioglaucine disodium salt dye retention in the mid intestine/vulva of *C. elegans*, each dot represents 10-25 animals. (e) Quantification of erioglaucine disodium salt dye retention in the lower intestine of *C. elegans*, each dot represents 10-25 animals.

**Table S1:**
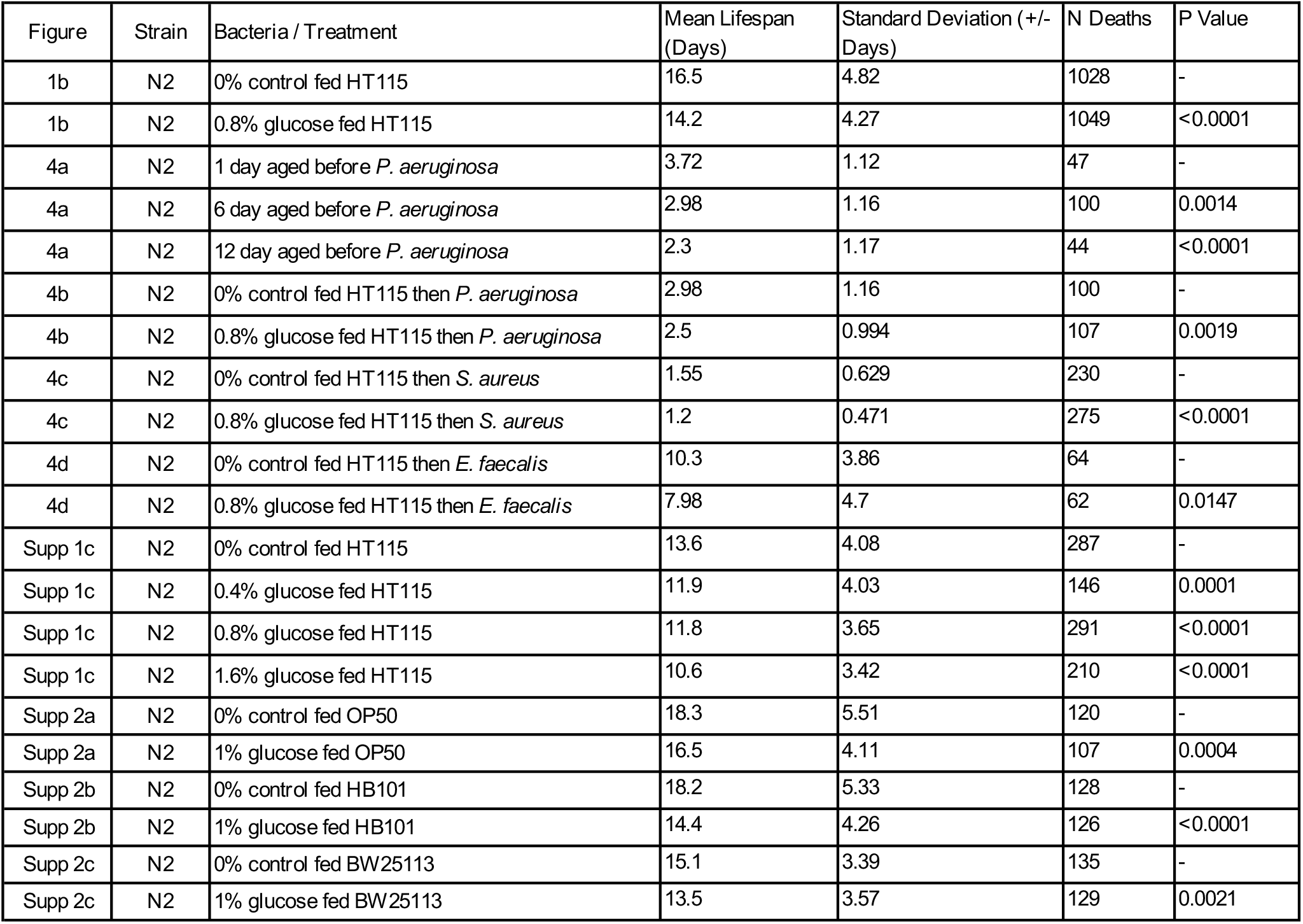
Lifespan Statistic.

| Figure | Strain | Bacteria / Treatment | Mean Lifespan (Days) | Standard Deviation (+/- Days) | N Deaths | P Value |
| --- | --- | --- | --- | --- | --- | --- |
| 1b | N2 | 0% control fed HT115 | 16.5 | 4.82 | 1028 | - |
| 1b | N2 | 0.8% glucose fed HT115 | 14.2 | 4.27 | 1049 | <0.0001 |
| 4a | N2 | 1 day aged before *P. aeruginosa* | 3.72 | 1.12 | 47 | - |
| 4a | N2 | 6 day aged before *P. aeruginosa* | 2.98 | 1.16 | 100 | 0.0014 |
| 4a | N2 | 12 day aged before *P. aeruginosa* | 2.3 | 1.17 | 44 | <0.0001 |
| 4b | N2 | 0% control fed HT115 then *P. aeruginosa* | 2.98 | 1.16 | 100 | - |
| 4b | N2 | 0.8% glucose fed HT115 then *P. aeruginosa* | 2.5 | 0.994 | 107 | 0.0019 |
| 4c | N2 | 0% control fed HT115 then *S. aureus* | 1.55 | 0.629 | 230 | - |
| 4c | N2 | 0.8% glucose fed HT115 then *S. aureus* | 1.2 | 0.471 | 275 | <0.0001 |
| 4d | N2 | 0% control fed HT115 then *E. faecalis* | 10.3 | 3.86 | 64 | - |
| 4d | N2 | 0.8% glucose fed HT115 then *E. faecalis* | 7.98 | 4.7 | 62 | 0.0147 |
| Supp 1c | N2 | 0% control fed HT115 | 13.6 | 4.08 | 287 | - |
| Supp 1c | N2 | 0.4% glucose fed HT115 | 11.9 | 4.03 | 146 | 0.0001 |
| Supp 1c | N2 | 0.8% glucose fed HT115 | 11.8 | 3.65 | 291 | <0.0001 |
| Supp 1c | N2 | 1.6% glucose fed HT115 | 10.6 | 3.42 | 210 | <0.0001 |
| Supp 2a | N2 | 0% control fed OP50 | 18.3 | 5.51 | 120 | - |
| Supp 2a | N2 | 1% glucose fed OP50 | 16.5 | 4.11 | 107 | 0.0004 |
| Supp 2b | N2 | 0% control fed HB101 | 18.2 | 5.33 | 128 | - |
| Supp 2b | N2 | 1% glucose fed HB101 | 14.4 | 4.26 | 126 | <0.0001 |
| Supp 2c | N2 | 0% control fed BW25113 | 15.1 | 3.39 | 135 | - |
| Supp 2c | N2 | 1% glucose fed BW25113 | 13.5 | 3.57 | 129 | 0.0021 |

**Table 2:**
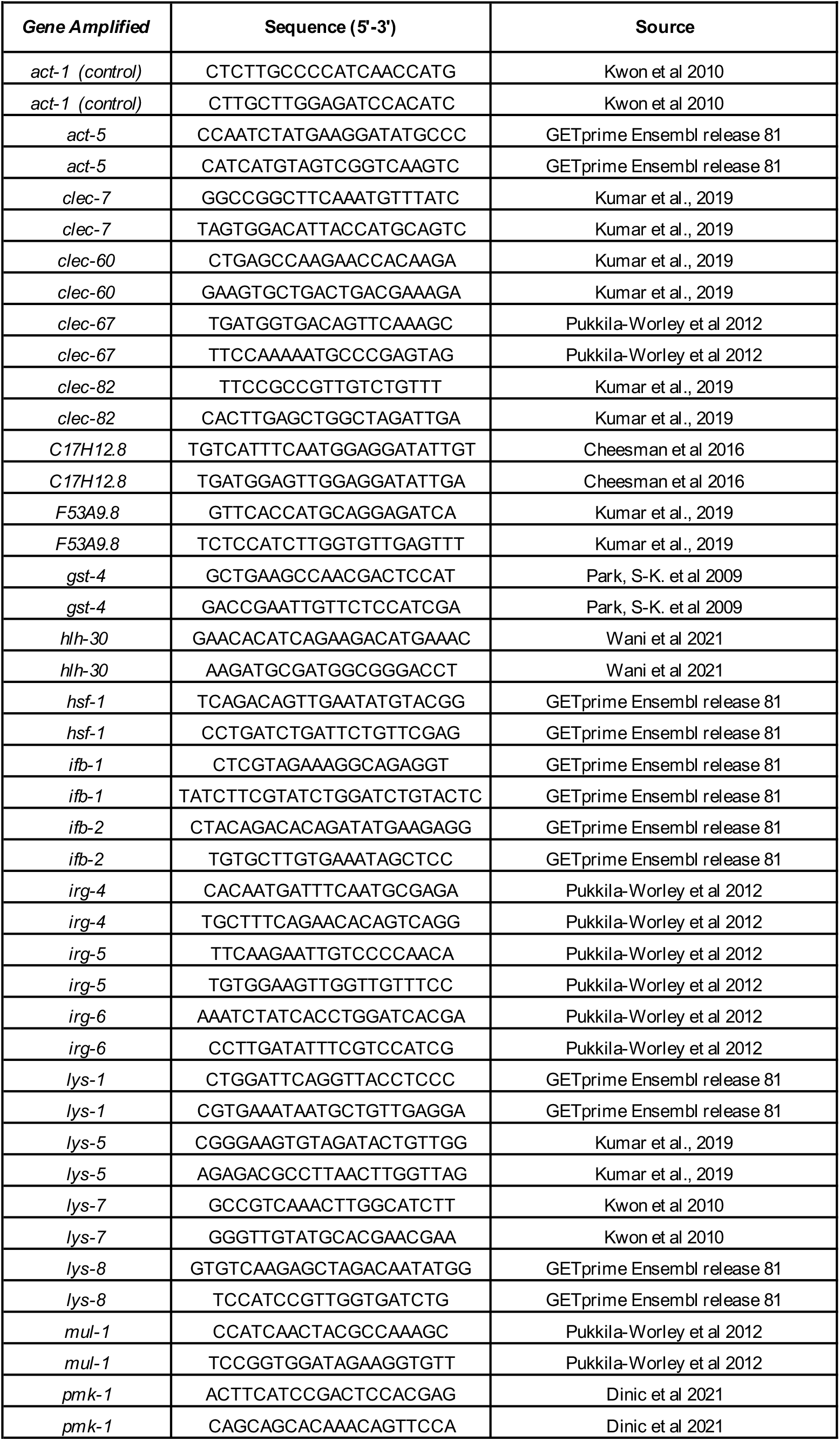
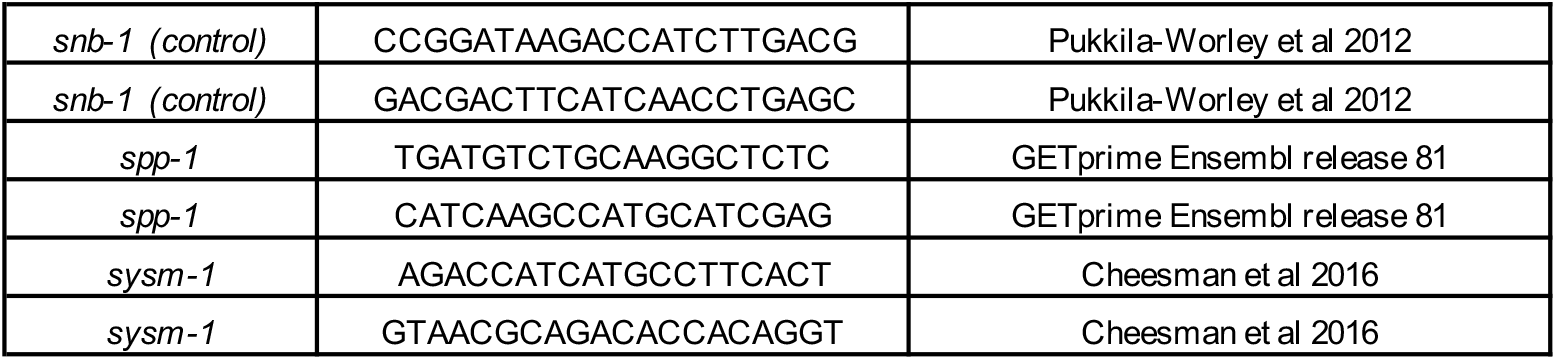
RTqPCR primer pair sequences.

## References

1 Di Rienzi, S. C. & Britton, R. A. Adaptation of the Gut Microbiota to Modern Dietary Sugars and Sweeteners. Adv Nutr 11, 616–629, doi:10.1093/advances/nmz118 (2020).

2 Khan, S. et al. Dietary simple sugars alter microbial ecology in the gut and promote colitis in mice. Sci Transl Med 12, doi:10.1126/scitranslmed.aay6218 (2020).

3 Do, M. H., Lee, E., Oh, M. J., Kim, Y. & Park, H. Y. High-Glucose or -Fructose Diet Cause Changes of the Gut Microbiota and Metabolic Disorders in Mice without Body Weight Change. Nutrients 10, doi:10.3390/nu10060761 (2018).

4 Choi, S. S. High glucose diets shorten lifespan of Caenorhabditis elegans via ectopic apoptosis induction. Nutr Res Pract 5, 214–218, doi:10.4162/nrp.2011.5.3.214 (2011).

5 Salim, C. & Rajini, P. S. Glucose feeding during development aggravates the toxicity of the organophosphorus insecticide Monocrotophos in the nematode, Caenorhabditis elegans. Physiol Behav 131, 142–148, doi:10.1016/j.physbeh.2014.04.022 (2014).

6 Liggett, M. R., Hoy, M. J., Mastroianni, M. & Mondoux, M. A. High-glucose diets have sex-specific effects on aging in C. elegans: toxic to hermaphrodites but beneficial to males. Aging (Albany NY) 7, 383–388, doi:10.18632/aging.100759 (2015).

7 Alcantar-Fernandez, J., Navarro, R. E., Salazar-Martinez, A. M., Perez-Andrade, M. E. & Miranda-Rios, J. Caenorhabditis elegans respond to high-glucose diets through a network of stress-responsive transcription factors. PLoS One 13, e0199888, doi:10.1371/journal.pone.0199888 (2018).

8 Alcantar-Fernandez, J. et al. High-glucose diets induce mitochondrial dysfunction in Caenorhabditis elegans. PLoS One 14, e0226652, doi:10.1371/journal.pone.0226652 (2019).

9 Kingsley, S. F. et al. Bacterial processing of glucose modulates C. elegans lifespan and healthspan. Sci Rep 11, 5931, doi:10.1038/s41598-021-85046-3 (2021).

10 Sabri, S., Nielsen, L. K. & Vickers, C. E. Molecular control of sucrose utilization in Escherichia coli W, an efficient sucrose-utilizing strain. Appl Environ Microbiol 79, 478–487, doi:10.1128/AEM.02544-12 (2013).

11 Raizen, D., Song, B. M., Trojanowski, N. & You, Y. J. Methods for measuring pharyngeal behaviors. WormBook, 1–13, doi:10.1895/wormbook.1.154.1 (2012).

12 Avery, L. The genetics of feeding in Caenorhabditis elegans. Genetics 133, 897–917 (1993).

13 Melo, J. A. & Ruvkun, G. Inactivation of conserved C. elegans genes engages pathogen- and xenobiotic-associated defenses. Cell 149, 452–466, doi:10.1016/j.cell.2012.02.050 (2012).

14 Hsu, A. L., Murphy, C. T. & Kenyon, C. Regulation of aging and age-related disease by DAF-16 and heat-shock factor. Science 300, 1142–1145, doi:10.1126/science.1083701 (2003).

15 Kumsta, C., Chang, J. T., Schmalz, J. & Hansen, M. Hormetic heat stress and HSF-1 induce autophagy to improve survival and proteostasis in C. elegans. Nat Commun 8, 14337, doi:10.1038/ncomms14337 (2017).

16 Garigan, D. et al. Genetic analysis of tissue aging in Caenorhabditis elegans: a role for heat-shock factor and bacterial proliferation. Genetics 161, 1101–1112 (2002).

17 Brunquell, J., Morris, S., Lu, Y., Cheng, F. & Westerheide, S. D. The genome-wide role of HSF-1 in the regulation of gene expression in Caenorhabditis elegans. BMC Genomics 17, 559, doi:10.1186/s12864-016-2837-5 (2016).

18 Baird, N. A. et al. HSF-1-mediated cytoskeletal integrity determines thermotolerance and life span. Science 346, 360–363, doi:10.1126/science.1253168 (2014).

19 Estes, K. A., Szumowski, S. C. & Troemel, E. R. Non-lytic, actin-based exit of intracellular parasites from C. elegans intestinal cells. PLoS Pathog 7, e1002227, doi:10.1371/journal.ppat.1002227 (2011).

20 Cabreiro, F. & Gems, D. Worms need microbes too: microbiota, health and aging in Caenorhabditis elegans. EMBO molecular medicine 5, 1300–1310, doi:10.1002/emmm.201100972 (2013).

21 Gelino, S. et al. Intestinal Autophagy Improves Healthspan and Longevity in C. elegans during Dietary Restriction. PLoS Genet 12, e1006135, doi:10.1371/journal.pgen.1006135 (2016).

22 Dukowicz, A. C., Lacy, B. E. & Levine, G. M. Small intestinal bacterial overgrowth: a comprehensive review. Gastroenterol Hepatol (N Y) 3, 112–122 (2007).

23 Chavez, V., Mohri-Shiomi, A., Maadani, A., Vega, L. A. & Garsin, D. A. Oxidative stress enzymes are required for DAF-16-mediated immunity due to generation of reactive oxygen species by Caenorhabditis elegans. Genetics 176, 1567–1577, doi:10.1534/genetics.107.072587 (2007).

24 Bolz, D. D., Tenor, J. L. & Aballay, A. A conserved PMK-1/p38 MAPK is required in caenorhabditis elegans tissue-specific immune response to Yersinia pestis infection. J Biol Chem 285, 10832–10840, doi:10.1074/jbc.M109.091629 (2010).

25 Irazoqui, J. E. et al. Distinct pathogenesis and host responses during infection of C. elegans by P. aeruginosa and S. aureus. PLoS Pathog 6, e1000982, doi:10.1371/journal.ppat.1000982 (2010).

26 Pukkila-Worley, R. & Ausubel, F. M. Immune defense mechanisms in the Caenorhabditis elegans intestinal epithelium. Curr Opin Immunol 24, 3–9, doi:10.1016/j.coi.2011.10.004 (2012).

27 Troemel, E. R. et al. p38 MAPK regulates expression of immune response genes and contributes to longevity in C. elegans. PLoS Genet 2, e183 (2006).

28 Peterson, N. D. & Pukkila-Worley, R. Caenorhabditis elegans in high-throughput screens for anti-infective compounds. Curr Opin Immunol 54, 59–65, doi:10.1016/j.coi.2018.06.003 (2018).

29 Anderson, S. M. et al. The fatty acid oleate is required for innate immune activation and pathogen defense in Caenorhabditis elegans. PLoS Pathog 15, e1007893, doi:10.1371/journal.ppat.1007893 (2019).

30 Cheesman, H. K. et al. Aberrant Activation of p38 MAP Kinase-Dependent Innate Immune Responses Is Toxic to Caenorhabditis elegans. G3 (Bethesda) 6, 541-549, doi:10.1534/g3.115.025650 (2016).

31 Stiernagle, T. Maintenance of C. elegans. WormBook, 1–11 (2006).

32 Garcia, A. M. et al. Glucose induces sensitivity to oxygen deprivation and modulates insulin/IGF-1 signaling and lipid biosynthesis in Caenorhabditis elegans. Genetics 200, 167–184, doi:10.1534/genetics.115.174631 (2015).

33 Margie, O., Palmer, C. & Chin-Sang, I. C. elegans chemotaxis assay. J Vis Exp, e50069, doi:10.3791/50069 (2013).

34 Ayala, F. R. et al. Culturing Bacteria from Caenorhabditis elegans Gut to Assess Colonization Proficiency. Bio Protoc 7, e2345, doi:10.21769/BioProtoc.2345 (2017).

35 Yoon, D. S., Lee, M. H. & Cha, D. S. Measurement of Intracellular ROS in Caenorhabditis elegans Using 2’,7’-Dichlorodihydrofluorescein Diacetate. Bio Protoc 8, doi:10.21769/BioProtoc.2774 (2018).

36 Pukkila-Worley, R. et al. Stimulation of host immune defenses by a small molecule protects C. elegans from bacterial infection. PLoS Genet 8, e1002733, doi:10.1371/journal.pgen.1002733 (2012).

37 Wani, K. A. et al. NHR-49/PPAR-alpha and HLH-30/TFEB cooperate for C. elegans host defense via a flavin-containing monooxygenase. Elife 10, doi:10.7554/eLife.62775 (2021).

## Supplemental References

1 Avery, L. & Shtonda, B. B. Food transport in the C. elegans pharynx. J Exp Biol 206, 2441–2457 (2003).

2 Houthoofd, K., Braeckman, B. P., Johnson, T. E. & Vanfleteren, J. R. Life extension via dietary restriction is independent of the Ins/IGF-1 signalling pathway in Caenorhabditis elegans. Exp Gerontol 38, 947–954 (2003).

3 Houthoofd, K. et al. No reduction of metabolic rate in food restricted Caenorhabditis elegans. Exp Gerontol 37, 1359–1369 (2002).

4 MacQueen, A. J. et al. ACT-5 is an essential Caenorhabditis elegans actin required for intestinal microvilli formation. Mol Biol Cell 16, 3247–3259, doi:10.1091/mbc.e04-12-1061 (2005).

